# KENT: Fast Pathogen Screening with NVIDIA Jetson Cluster in Resource-Constrained Areas

**DOI:** 10.64898/2026.09.28.754999

**Authors:** Arda Icoz, Ricardo Roman-Brenes, Zülal Bingöl, Can Alkan

## Abstract

Metagenomic sequencing enables broad pathogen detection without requiring prior assumptions about the organisms present, but the memory and computational demands of classifying large sequencing datasets against reference databases often lead analysis pipelines to rely on desktop or server-grade computing platforms. This limits practical use in disaster-stricken and other resource-constrained environments, where portable sequencing may be available but reliable access to cloud or laboratory infrastructure is not. We present KENT, a portable edge-computing framework for metagenomic pathogen screening based on a cluster of NVIDIA Jetson Nano modules. KENT adapts CuCLARK-l for low-memory embedded execution and extends it with a custom software stack for distributed read classification, abundance estimation, split-run abundance merging, and report generation. Using an MPI-based controller/responder architecture, KENT supports local, offline analysis with sample-level parallelism across multiple nodes.

We evaluate KENT using long read wastewater metagenomic samples with a custom pathogen database and the CAMI2 Marine long read dataset. In the wastewater setting, KENT achieves an average speedup of 3.1*×* over the CPU-based CLARK-l baseline while maintaining close agreement in classified read abundance estimates. On CAMI2 Marine, KENT achieves an average speedup of 2.68*×* and similarly preserves CLARK-l abundance profiles. Evaluation against the CAMI2 long read taxonomic ground truth yields a mean species-level precision of 58.55%, recall of 93.36%, F1 score of 71.74%, and L1 abundance distance of 0.731, closely matching CLARK-l. Among ground truth species represented in the reference database, KENT detects 99.86% on average. A controlled database-size experiment further shows little change in classification throughput across the tested database configurations, while large sequencing inputs remain constrained by the Jetson Nano’s unified memory capacity. Overall, KENT demonstrates that GPU-accelerated metagenomic classification can be adapted to low-memory embedded hardware while preserving the biological profiling behavior of the CLARK-l baseline, supporting portable and offline pathogen screening in resource-constrained settings.

**Availability:** KENT is available at https://github.com/BilkentCompGen/KENT

## 1 Introduction

Natural disasters frequently disrupt water, sanitation, electricity, and communication infrastructure, creating conditions in which infectious disease outbreaks, such as cholera and dysentery [1], can emerge rapidly. In such environments, timely pathogen screening is important not only for clinical response but also for environmental surveillance, especially when water and food sources become contaminated [2]. Conventional laboratory workflows are ill-suited to these conditions: culture-based methods are often too slow, while targeted molecular assays, such as PCR, require prior assumptions about which organisms to test. Read classification from metagenomic sequencing data offers a more flexible alternative, enabling broad, untargeted detection directly from mixed biological samples [3, 4]. Portable sequencers (e.g., Oxford Nanopore Technologies (ONT) MinION) have enabled field-side data generation [5], but the downstream analysis step still typically depends on powerful workstations, high-performance computing, or cloud connectivity. This dependency creates a major gap between portable sequencing and truly portable pathogen screening in disaster-stricken, resource-limited settings.

Edge computing is an attractive deployment model for metagenomic analysis in the field. Instead of transferring large sequencing datasets to remote servers, edge systems process reads locally, reducing reliance on network connectivity and enabling operation even when internet access is unavailable or unstable. This is particularly crucial in post-disaster environments, where infrastructure failure can make cloud-based analysis impractical. Edge devices provide portability and low power consumption by operating under much tighter compute and memory budgets than conventional desktop or server hardware. However, the central challenge is not simply to run metagenomic classification outside the laboratory, but to do so reliably on severely resource-constrained hardware. Embedded devices, such as the NVIDIA Jetson Nano modules, offer a portable, highly parallel computing platform that can facilitate large-scale metagenomic analysis compared to CPU-only pipelines. Their main computational advantage is an onboard GPU that supports CUDA, in addition to an ARM-based CPU. A GPU-oriented design, therefore, provides a practical way to partially compensate for the reduced capabilities of embedded edge hardware, especially for read classification workloads that can benefit from large-scale parallelism. Therefore, GPU acceleration is not merely an optimization; it is one of the main enablers that makes field-deployable pathogen screening computationally plausible.

Several GPU-accelerated metagenomic classifiers have demonstrated that read classification can be substantially accelerated on hardware. For example, GPMeta [6] is a recent GPU-accelerated framework for ultrarapid pathogen identification from metagenomic sequences, while MetaCache-GPU [7] emphasizes high-performance database construction and large-scale classification. Both are modern GPU-based tools for metagenomic classification, but they are designed for more powerful platforms than the ARM-based Jetson Nano used here We address a different deployment setting: low-power, memory-constrained embedded edge devices operating with an older but more affordable CUDA stack and without reliance on external infrastructure. Other metagenomic workflows, such as MetaPORE [8], MAIRA [9], and NanoMeta Live [10], emphasize real-time analysis, pathogen detection, visualization, or user-facing interpretation during sequencing. In contrast, our focus is on offline, self-contained classification on a low-power GPU cluster designed to operate under strict memory, power, and internet connectivity constraints.

Under these constraints, CuCLARK [11] emerges as the most practical GPU-based software for our system. CuCLARK is a CUDA-enabled extension of CLARK [12] that performs read-level metagenomic classification using reduced *k*-mers. Crucially for this work, CuCLARK provides a light mode, CuCLARK-l (like CLARK-l [12]), that reduces memory usage during classification, directly addressing the Jetson Nano platform’s low memory limit. For these reasons, CuCLARK-l is the most practical candidate for adaptation to our deployment target while still supporting memory-reduced execution.

In this paper, we present KENT, a portable edge-computing framework for metagenomic pathogen screening built for a cluster of NVIDIA Jetson Nano modules. Our design bridges the gap between portable sequencing and portable analysis by enabling local, offline read classification in environments with limited power, memory, and connectivity. KENT delivers an end-to-end pathogen screening pipeline for the healthcare personnel in the field to easily execute with minimal technical intervention. To achieve this, we adapt CuCLARK-l for embedded execution, integrate it with a custom workflow for database construction, classification, abundance estimation, split-run abundance merging, and report generation, and extend the pipeline to a distributed controller/responder architecture using Message Passing Interface (MPI). The main contributions of this work are: (1) a deployable Jetson Nano GPU–based edge frame-work for offline pathogen screening; (2) an embedded adaptation of CuCLARK-l that supports execution under strict memory and runtime constraints; (3) a distributed software stack for end-to-end classification and reporting; and (4) an evaluation showing that the system preserves close agreement with CPU-based CLARK-l abundance estimates while operating on low-power edge hardware.

## 2 Methods

Here, we present a GPU-accelerated metagenomic read classification tool for portable deployment in resource-constrained environments. We focus on transforming CuCLARK-l into a field-deployable pathogen screening framework that can operate on a low-power embedded cluster without relying on cloud infrastructure. Our proposed system combines porting CuCLARK-l source code with a custom orchestration layer and a distributed execution model tailored to the hardware constraints of NVIDIA Jetson Nano modules. This design enables localized pathogen screening in disaster-stricken settings where portability, low power consumption, and offline operation are essential.

### 2.1 Cluster Topology and Software Architecture

The physical platform consists of several NVIDIA Jetson Nano modules organized in a controller/responder topology, using 11 modules in this study. One node serves as the controller and is responsible for coordination, configuration broadcast, and result aggregation through MPI. The remaining nodes serve as responders and execute classification tasks on locally stored sequencing datasets. The number of nodes can be scaled up or down depending on available equipment.

The software architecture is organized into three layers (Figure 1):

**Figure 1:**
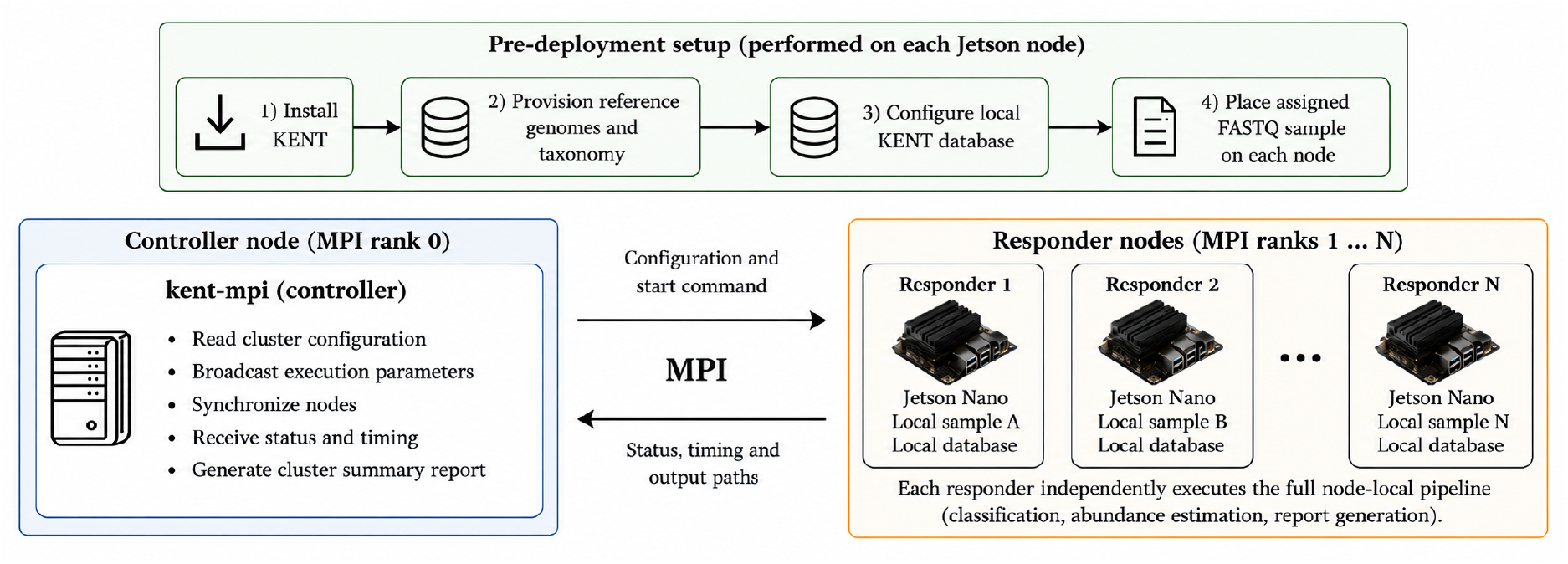
KENT controller–responder architecture. Each Jetson node is pre-provisioned with KENT, a local reference database, and an assigned FASTQ sample. The controller uses kent-mpi to broadcast parameters, synchronize execution, and collect status, timing, and output metadata. Responders independently perform classification, abundance estimation, and report generation, providing sample-level parallelism with limited network communication.

- **Modified CuCLARK-l backend:** CuCLARK-l serves as the core GPU-accelerated read classifier. Its source code and execution behavior were adapted to improve compatibility with the Jetson Nano GPU and to support reliable operation under strict memory constraints.
- **Node-local orchestration layer (**kent): kent is a custom C++ control module responsible for database verification, target database construction, classification execution, abundance estimation, abundance merging for fragmented runs, and report generation on each node.
- **Distributed coordination layer (**kent_mpi): kent_mpi extends the single-node workflow to the cluster level by using MPI to broadcast execution parameters, synchronize responder nodes, and collect execution outputs.

CuCLARK-l is the main classification engine, while the proposed framework extends it through embedded hardware adaptation, execution control, distributed coordination, and post-processing.

### 2.2 Hardware and Deployment Setup

The experimental platform consisted of 11 NVIDIA Jetson Nano modules connected to form a local edge computing cluster. Each Jetson Nano is equipped with an NVIDIA Tegra X1 system-on-chip, including a Maxwell-generation CUDA-capable GPU and an ARM-based CPU, and provides 2 GB of unified system memory shared between CPU and GPU workloads. The modified CuCLARK-l backend was compiled for CUDA compute capability sm_53, corresponding to the Jetson Nano GPU architecture, and the experimental software environment used CUDA 10.2.

All Jetson Nano nodes were operated in the NVIDIA MAXN power mode (NVPModel mode 0), corresponding to a nominal 10 W module-level power budget. The same power configuration was used on the controller and all ten responder nodes throughout the experiments. Thus, when all 11 modules are active, the cluster represents a nominal aggregate module-level power budget of 110 W. This value refers to the configured Jetson module power mode rather than measured wall-power consumption; carrier-board, storage, networking, cooling, and other peripheral power are not included.

The nodes were connected through a local 10/100 Mbps Ethernet network. MPI communication was restricted to the eth0 interface, while sequencing inputs and reference databases were provisioned locally on the participating nodes before classification. Consequently, large FASTQ inputs and reference databases did not need to be repeatedly transferred through the network during analysis. Passwordless SSH was used to launch MPI processes between nodes, while the classification workflow itself communicated via MPI. This design limits network traffic during classification and allows operation with modest networking infrastructure.

The Jetson Nano platform imposes several constraints that directly influence KENT’s execution strategy. Most importantly, GPU workloads share the same limited 2 GB memory pool rather than using separate system and GPU memory. Database storage, input batches, intermediate classification results, and operating-system processes therefore compete for a small common memory budget. This limitation motivated the use of CuCLARK-l, which follows CLARK-l’s strategy of constructing a sparse database from sampled, non-overlapping reference k-mers, thereby reducing the memory required for the database index. To further reduce memory requirements per input batch, KENT uses more batches for memory-intensive workloads. For inputs that still exceed the available memory budget, KENT relies on externally partitioned FASTQ files followed by abundance-level merging.

KENT uses the Jetson cluster primarily for sample-level throughput: separate sequencing samples are assigned to different nodes and processed concurrently. The current implementation does not distribute the reads of a single sample across multiple Jetson GPUs. Therefore, adding responder nodes increases the number of independent samples that can be processed concurrently, but does not directly reduce the classification latency of an individual sample. These properties make the platform suitable for offline, multi-sample field screening while leaving memory capacity as the principal limitation for large reference databases and sequencing inputs.

### 2.3 Usage of CuCLARK-l in KENT

KENT retains the core CuCLARK-l workflow, including discriminative k-mer database construction, FASTA/FASTQ processing, configurable input batching, GPU-based classification, and CLARK-compatible classification output. KENT adapts this classifier to the Jetson Nano modules, considering the resource constraints, and integrates it into a complete edge-computing workflow.

Around the modified CuCLARK-l backend, KENT adds a node-local orchestration layer for database verification and configuration, classification execution, abundance estimation, split-run abundance merging, and report generation. It further adds an MPI-based coordination layer that enables independent samples to be processed concurrently across multiple Jetson nodes. Figure 2 summarizes the relationship between the original CuCLARK workflow and the additional adaptations and workflow components introduced by KENT.

**Figure 2:**
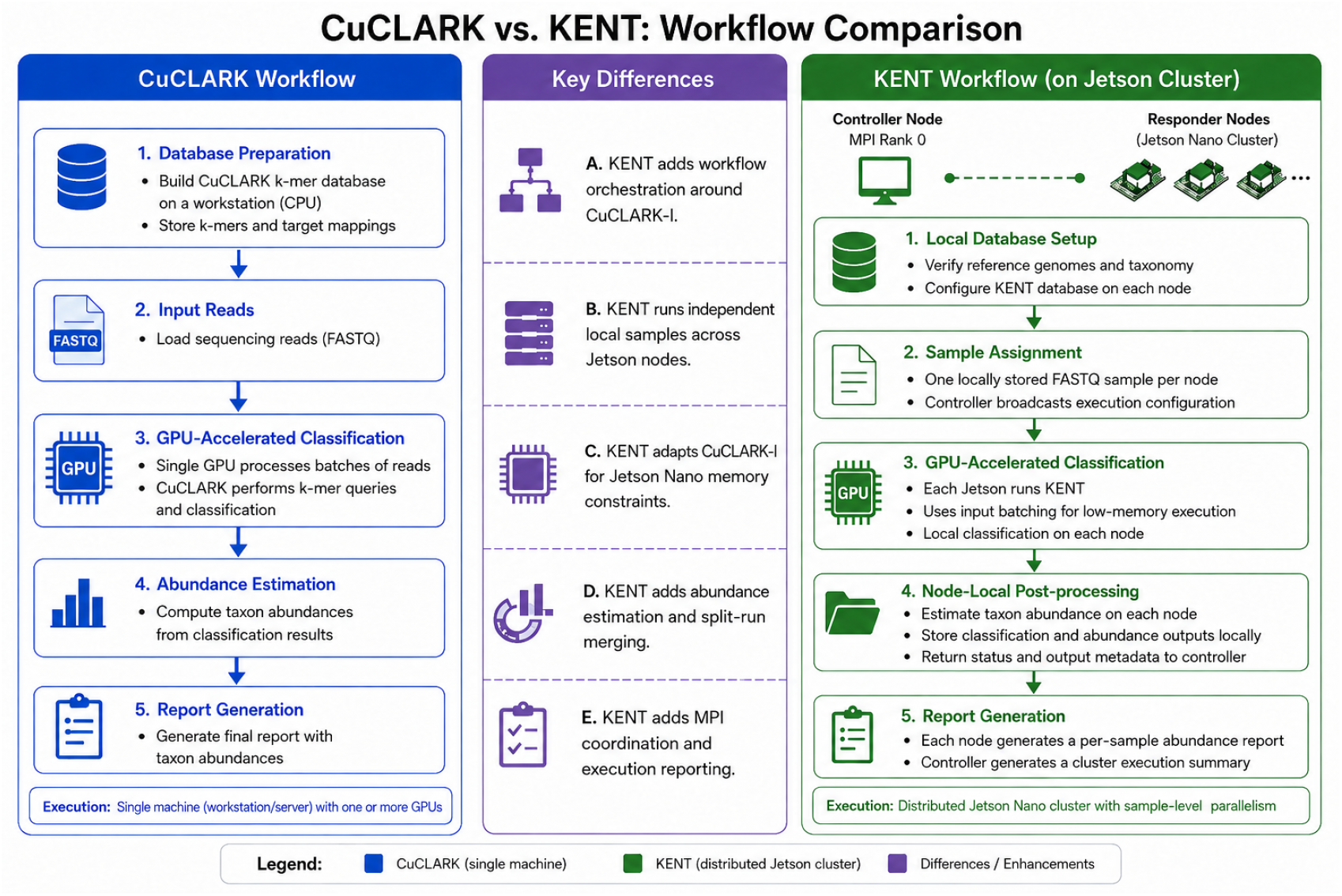
Comparison of the original CuCLARK workflow and the KENT workflow. KENT retains the CuCLARK-l classification core while adapting its execution for the Jetson Nano memory environment and adding node-local orchestration, abundance processing, split-run merging, report generation, and MPI-based sample-level coordination across multiple Jetson modules.

### 2.4 Reference Database Construction

KENT uses a custom reference database constructed specifically for resource-constrained pathogen screening. Rather than relying on a broad general-purpose reference space, the workflow is designed to build a targeted database that can be executed efficiently on low-memory embedded hardware. This design reduces unnecessary computational overhead and makes the classification pipeline more suitable for portable deployment. In our experiments, we consulted with microbiologists at Bilkent City Hospital in Ankara, Turkey, for the pathogens that are most likely to be encountered; this custom database can be changed depending on the needs and the specific ecological properties of the locations where the tool will be deployed. The list of pathogens included in this database is provided in Appendix A1 of the supplementary material.

The kent control layer prepares the database as a pre-deployment operation. Before configuring the database, kent verifies the local environment to ensure that the required reference genome files and NCBI taxonomy resources are present and correctly organized. After validation, the selected reference set is prepared in the database format required by CuCLARK-l. This procedure allows the participating Jetson nodes to be provisioned with a consistent reference database before distributed classification begins, reducing manual configuration errors and improving reproducibility across devices.

### 2.5 Database Size Sensitivity Experiment

To evaluate the effect of reference database size independently of sample input size, we constructed three nested subsets of the CAMI2 Marine reference database. Starting from the 501 unique taxonomy IDs (i.e., TaxIDs) represented in the complete database, the TaxIDs were placed in a reproducibly randomized order using a fixed random seed of 42. Nested target sets containing the first 100, 300, and 501 TaxIDs were then selected. All FASTA files associated with the selected TaxIDs were included in the corresponding reference subset, and each subset was processed independently using the standard KENT database preparation procedure. After taxonomy mapping and removal of unidentified references, the resulting databases contained 96, 289, and 501 unique mapped TaxIDs, respectively.

For the performance comparison, all three databases were evaluated on the same NVIDIA Jetson Nano node using the same CAMI2 Marine fragment sample_0. The input contained 822,406 reads and corresponded to one of two approximately equal fragments of the original sample. CuCLARK-l was configured with 128 input batches for all runs. No other classification parameters were changed between database configurations. For each database, we measured the compiled database size on disk, the database size reported by CuCLARK-l when loaded in RAM, the number of database-processing loops required, total classification runtime, and classification throughput. This controlled design keeps the input and execution configuration fixed while varying the size of the reference database.

### 2.6 Ground-Truth Taxonomic Evaluation

In addition to measuring agreement between KENT and the CLARK-l baseline, we evaluated both methods against the CAMI2 Marine long-read PacBio taxonomic ground truth. Each of the ten long read samples was matched to its corresponding CAMI2 taxonomic profile. Evaluation was performed at the species level and was restricted to bacterial and archaeal species, since these represent the cellular organisms targeted by the Marine reference database. Plasmid, viral, and unidentified ground-truth entries were excluded from this species-level evaluation.

A species was considered detected when the classifier assigned at least one read to its TaxID. For each sample, we calculated precision, recall, and F1 score using the set of species present in the CAMI2 ground truth and the set of species reported by the classifier. Precision was defined as the fraction of detected species that were present in the ground truth, recall as the fraction of ground-truth species that were detected, and F1 as the harmonic mean of precision and recall. To evaluate abundance estimation, we additionally calculated the L1 distance between the predicted and ground-truth species-abundance distributions. Ground truth abundances were normalized over the evaluated bacterial and archaeal species, and classifier abundances were normalized over the corresponding classified species before comparison. The L1 distance was calculated as

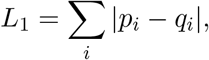

where *p*_*i*_ and *q*_*i*_ denote the normalized ground-truth and predicted abundance of species *i*, respectively. An L1 value of 0 indicates identical abundance distributions, while larger values indicate greater disagreement, with a maximum possible value of 2 for normalized distributions. Because the custom Marine reference database does not include all bacterial and archaeal species present in the CAMI2 ground truth, we also performed a secondary, database-aware analysis. In this analysis, recall was calculated only over ground-truth species whose TaxIDs were represented in the KENT reference database. This additional metric separates classification failures from species that could not be detected because their reference genomes were absent from the database. The conventional, database-independent precision, recall, F1, and L1 metrics were retained as the primary evaluation.

### 2.7 Source-Level Adaptation of CuCLARK-l for Embedded Execution

One of the main methodological contributions of KENT is the adaptation of CuCLARK-l to operate within the restricted compute and memory environment of the NVIDIA Jetson Nano. Rather than modifying the underlying discriminative k-mer classification procedure, the changes target platform compatibility, memory initialization, and workload configuration.

At the build level, the CUDA compilation target was changed from sm_30 in the original CuCLARK implementation to sm_53, corresponding to the Maxwell-generation GPU used by the Jetson Nano. The runtime memory assumptions were also relaxed for the embedded platform. The minimum free GPU-memory check was reduced from 1 GB to 200 MB, and the memory reserved by the CuCLARK-l configuration was reduced from 300 MB to 150 MB. These changes allow the classifier to initialize and execute under the substantially smaller memory budget available on the Jetson Nano. The main implementation-level differences are summarized in Appendix Table A17.

At the execution level, kent invokes the light-memory CuCLARK-l variant and exposes CuCLARK’s existing input-batching mechanism. The -b parameter specifies the number of batches into which the sequencing input is divided; it does not specify a fixed number of reads per batch. Increasing the number of input batches, therefore, reduces the amount of sequence data and intermediate storage required per batch. KENT uses this parameter to adapt the workload to the available memory, with larger batch sizes used for more demanding combinations of sample and database sizes.

These modifications preserve the CuCLARK-l classification procedure while adapting its compilation, initialization, and workload configuration to the Jetson Nano environment. The CLARK-l CPU baseline does not provide an equivalent user-configurable input-batching parameter and did not exhibit the same memory-related execution failures in our experiments.

### 2.8 Sample-Level Parallelism and Distributed MPI Workflow

The distributed execution model is based on sample-level parallelism. Each responder node is pre-provisioned with one complete local read dataset, typically in FASTQ format, together with a local copy of the target database. This design avoids repeated network transmissions of large sequencing datasets and reduces reliance on available bandwidth during field operations.

At runtime, the controller broadcasts the serialized execution configuration to the participating MPI ranks and synchronizes the nodes before classification begins. Each responder identifies the input assigned to its hostname and independently executes the complete node-local workflow. This workflow invokes *kent*, runs the modified CuCLARK-l backend, stores the classification output, and performs abundance estimation.

After processing, each responder sends a compact execution record to the controller. This record contains execution metadata, including the node hostname, success or failure status, elapsed time, error information, and the paths of the generated classification and abundance files. The complete FASTQ inputs and classification result files are not transferred through MPI. Consequently, the cluster implements sample-level parallelism rather than distributing the reads of an individual sample across multiple Jetson modules, limiting network communication during classification. The node-local execution procedure is illustrated in Figure 3.

**Figure 3:**
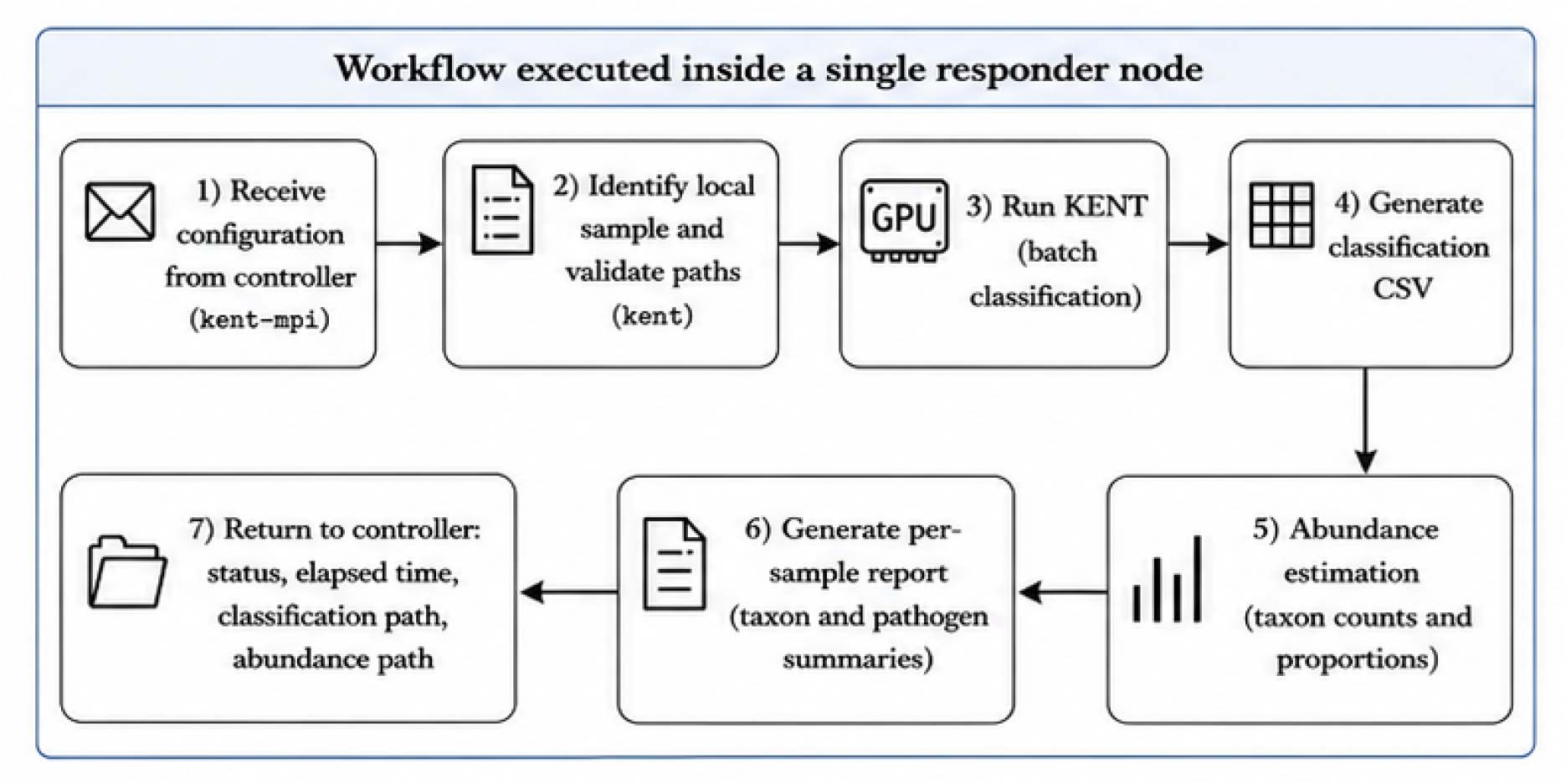
Node-local KENT workflow. After receiving the MPI configuration, a responder identifies and validates its locally assigned sequencing sample, invokes the modified CuCLARK-l backend through *kent*, stores the classification output, performs abundance estimation, generates a per-sample report, and returns execution status, elapsed time, and output-path metadata to the controller.

### 2.9 Fallback Handling for Oversized Samples

Although increasing the number of input batches stabilizes classification for many samples, exceptionally large inputs can still exceed the Jetson Nano’s available unified memory budget. In these cases, the input FASTQ file is externally partitioned into smaller fragments before classification. Each fragment is then processed as an independent KENT run, producing its own classification and abundance output. The resulting fragment-level abundance files are subsequently merged by KENT to reconstruct a final abundance profile for the original sample.

This fragmentation procedure provides a fallback path for samples that cannot be processed reliably in a single run. Input partitioning itself is performed before invoking KENT, whereas the subsequent abundance merging is implemented within the kent workflow.

### 2.10 Post-Processing, Abundance Estimation, and Merging of Split Runs

After classification, *kent* invokes the CLARK-compatible abundance estimation procedure on the CuCLARK-l classification output. For each detected taxon, the resulting abundance file reports the raw assigned read count, the proportion among all reads, and the proportion among classified reads. Assignments reported as UNKNOWN are handled separately so that classified read proportions are calculated using only reads assigned to known taxa.

For fragmented samples, *kent* implements an explicit abundance merging procedure. Each fragment-level abundance file is parsed into entries keyed by taxonomic identifier, and counts corresponding to the same taxon are summed across all fragments. After aggregation, the total number of reads and the number of classified reads are recomputed, and abundance proportions are recalculated from the merged counts. UNKNOWN assignments are preserved separately and excluded from the denominator used for classified read proportions.

This procedure allows independently classified fragments of the same original sample to be recombined without averaging already normalized abundance values. Instead, the final proportions are recalculated from the aggregated raw counts. The complete split-run procedure is illustrated in Figure 4.

**Figure 4:**
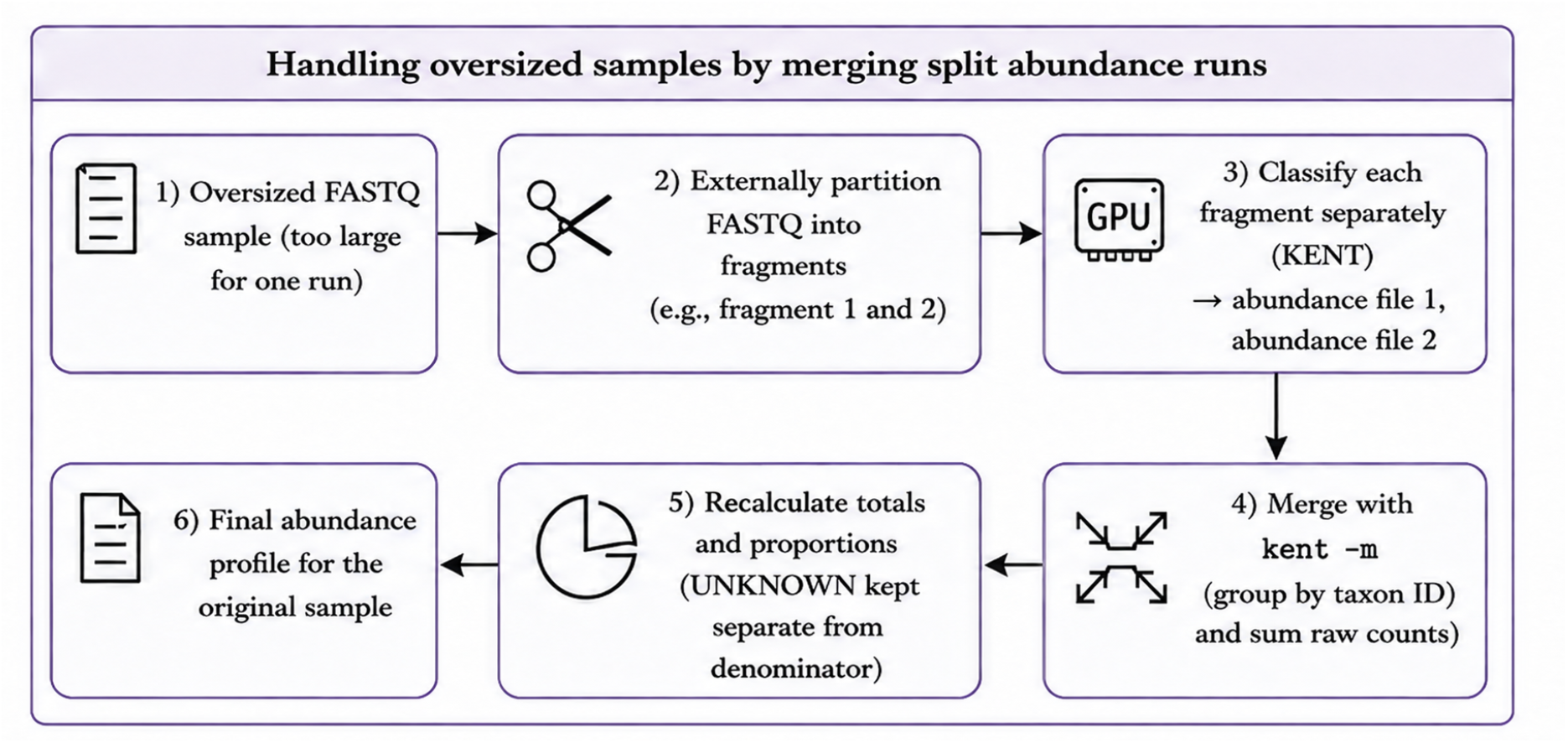
Split-run handling for oversized FASTQ samples. Oversized inputs are externally divided into fragments, classified independently, and merged by summing taxon-level counts and recalculating abundance proportions. UNKNOWN assignments are preserved separately when classified-read proportions are computed.

## 3 Results & Discussion

### 3.1 Wastewater Dataset

The first evaluation was conducted on a wastewater metagenomic dataset [13] because water quality assessment is directly relevant to disaster-stricken environments. The dataset consists of 11 ONT long read FASTQ samples from a public wastewater metagenomics study. Sample sizes vary substantially, ranging from 378,859 to 839,634 reads and from approximately 2.44 to 5.40 Gbp in total sequence length.

For this experiment, a targeted, custom pathogen database was constructed with support from medical experts, as previously described in Section 2.4. The database contains 22 reference sequences. The complete list is available in the Appendix Table A1. The source reference collection occupied approximately 118 MB on disk. After CuCLARK-l database construction, the compiled canonical database occupied approximately 62 MB on disk and was reported by CuCLARK-l as 0.237 GB when loaded in RAM. The database was processed in a single CuCLARK-l database loop.

This wastewater experiment was executed using the default *kent* classification configuration with 32 input batches. However, several high-density samples, such as those containing total sequence lengths exceeding 4.4 Gbp, could not be processed safely in a single pass within the Jetson Nano module’s memory limit. In these cases, the input FASTQ files were externally partitioned into two fragments before classification, and the resulting abundance files were subsequently merged using the split-run abundance-merging procedure described in the methodology.

Table 2 summarizes the agreement and performance of KENT relative to the CLARK-l baseline on both datasets. Database size denotes the in-memory database size reported by CuCLARK-l at runtime. Agreement is measured using the mean absolute difference (MAD) in classified read abundance proportions, with lower values indicating closer agreement. Speedup is measured relative to CLARK-l, with higher values indicating faster execution. For the wastewater dataset, classification consistency was assessed by comparing the abundance outputs from the modified CuCLARK-l pipeline with those from CLARK-l, using the same reference database and a Jetson-based environment. The comparison used the proportion of classified reads assigned to each taxon. Across the 11 wastewater samples, agreement between the two implementations remained strong. The mean absolute difference in taxon-level classified read proportions was approximately 0.12% when averaged over all samples, with per-sample mean absolute differences ranging from about 0.07% to 0.19%. The dominant taxon was preserved across all 11 samples: in 9 samples, both implementations identified *Plasmodium falciparum* as the most abundant taxon, whereas in the remaining 2 samples, both implementations identified *Escherichia coli* as the dominant taxon. These results indicate that KENT preserves the abundance profile of the CLARK-l baseline with only minor deviations at the taxon level.

**Table 1.** Hardware and deployment configuration used for KENT evaluation.

| Component | Configuration |
| --- | --- |
| Compute platform | NVIDIA Jetson Nano |
| Number of nodes | 11 (1 Controller, 10 Responders) |
| System-on-chip | NVIDIA Tegra X1 |
| GPU architecture | Maxwell, CUDA compute capability 5.3 |
| CUDA Cores | 128 on one node |
| Memory | 2 GB unified memory per node |
| CUDA version | 10.2 |
| Network | Local Ethernet network |
| MPI interface | `eth0` |
| Execution model | One locally provisioned sample per responder |
| Reference data | Local copy on each participating node |
| Internet requirement during classification | None |
| Power mode | MAXN |

**Table 2.** Agreement and performance of KENT compared with the CLARK-l baseline on two datasets.

| Dataset | Database in memory (GB) | Sample Length (Gbp) |  |  | MAD* (%) |  | Speedup |
| --- | --- | --- | --- | --- | --- | --- | --- |
|  |  | Avg. | Min. | Max. | Taxon Level | Per Sample |  |
| Wastewater [13] | 0.237 | 3.5 | 2.4 | 5.4 | 0.12 | 0.07 – 0.19 | 3.10× |
| CAMI2 Marine [14] | 0.353 | 4.8 | 4.7 | 4.9 | 0.01 | 0.01 | 2.68× |

\* MAD: Mean Absolute Difference (lower is better)

In terms of runtime, KENT shows a clear advantage for the GPU-accelerated implementation. When both tools were executed on Jetson modules, KENT consistently outperformed the CLARK-l baseline across all 11 samples (detailed runtimes are provided in Appendix A4 and A5). Averaged over the matched runs, KENT completed classification in roughly one-third of the time required by CLARK-l, corresponding to an average speedup of approximately 3.1×. The magnitude of this improvement varied across samples, but the trend was consistent, even in the fragmented high-density runs. This result is important because it shows that GPU acceleration is not merely a portability compromise on edge hardware; on the target embedded platform, it provides a substantial throughput advantage while maintaining abundance estimates close to those produced by the CLARK-l baseline.

### 3.2 CAMI2 Marine Long Read Dataset

A second evaluation was conducted using the PacBio long read portion of the CAMI2 Marine dataset [14] to test the system on a larger, more diverse reference space. Unlike the wastewater experiment, which relies on a compact targeted pathogen panel, this setting uses the genome set provided with the dataset itself for reference database construction. The complete source reference directory occupied approximately 2.8 GB on disk. During KENT database preparation, 618 FASTA files were successfully retained in the target definition, corresponding to 501 unique taxonomic targets. The resulting compiled canonical CuCLARK-l database occupied approximately 172 MB on disk and was reported as 0.353 GB when loaded in RAM. Like the wastewater database, it was processed in a single CuCLARK-l database loop.

The combination of the larger reference space and the approximately 4.7 to 4.9 Gbp input samples imposed greater memory pressure on the Jetson Nano platform. Under the default execution configuration, classification terminated with a Killed message, indicating that the workload exceeded the available memory budget. To stabilize execution, the number of input batches was increased to 128. In addition, the input read files were partitioned into smaller fragments before classification, and the fragment-level outputs were subsequently merged to reconstruct final abundance estimates for each sample. Notably, this behavior was observed consistently across the CAMI2 Marine dataset. The complete samples ranged from approximately 4.70 to 4.91 Gbp and could not be processed reliably as single inputs on the Jetson Nano under the tested configuration. After each sample was divided into two approximately equal fragments, corresponding to roughly 2.35 to 2.46 Gbp per fragment, classification was completed successfully. These observations provide an empirical operating range for the tested hardware and software configuration: inputs near 4.7 Gbp or larger required fragmentation, whereas fragments of approximately 2.4 Gbp were processed successfully. The exact transition point was not determined and may depend on database size, read-length distribution, batch count, and the amount of system memory available at runtime.

The CAMI2 Marine experiment was designed to test whether the proposed system could remain consistent with the CLARK-l baseline when the reference database size moved well beyond the compact disaster-screening scenario used in the wastewater experiment. Table 2 summarizes the agreement and performance of KENT relative to the CLARK-l baseline. Comparative analysis against the CLARK-l baseline showed very close agreement in the resulting abundance profiles. Across the 10 comparison files, the mean absolute difference in taxon-level classified read proportion was 0.01% when averaged over samples, with per-sample mean absolute differences ranging between 0.0052% and 0.0070%. The dominant classified organism was preserved across all 10 samples, with both implementations identifying *Alteromonas mediterranea* as the most abundant taxon. These results indicate that KENT remained highly consistent with the CLARK-l baseline even under a substantially larger reference space.

Runtime behavior again favored the GPU-accelerated implementation on the target embedded platform. For the CAMI2 samples with matched runtime logs available in both implementations, the KENT consistently outperformed CLARK-l on Jetson modules. The observed speedup ranged from approximately 1.65*×* to 4.69*×*, with an average speedup of about 2.68× across the matched runs. This result is particularly important because the CAMI2 setting required a much larger reference database, a more aggressive batching strategy, and routine split-run processing to remain within the memory limits of the Jetson Nano modules. Despite these constraints, the GPU-based pipeline still provided a clear throughput advantage while preserving abundance estimates close to those of the CLARK-l baseline.

### 3.3 Ground-Truth Taxonomic Evaluation

We further evaluated KENT and the CLARK-l baseline against the CAMI2 Marine long-read taxonomic ground truth. Table 3 summarizes the species-level results across all ten samples. The evaluation was restricted to bacterial and archaeal species as described in Section 2.6.

**Table 3.** Species-level evaluation against the CAMI2 Marine long-read ground truth. Values are the mean *±* standard deviation across the ten samples. Precision, recall, and F1 are reported as percentages. Lower L1 values indicate closer agreement with the ground-truth abundance distribution.

| Method | Precision (%) | Recall (%) | F1 (%) | L1 |
| --- | --- | --- | --- | --- |
| CLARK-l | 58.33 ± 7.12 | 93.42 ± 0.93 | 71.59 ± 5.25 | 0.729 ± 0.060 |
| KENT | 58.55 ± 7.16 | 93.36 ± 0.97 | 71.74 ± 5.24 | 0.731 ± 0.061 |

**Table 4.** Effect of reference database size on KENT classification performance. All runs used the same 822,406-read CAMI2 Marine FASTQ fragment and 128 input batches on the same Jetson Nano.

| Mapped TaxIDs | Compiled DB (MB) | DB in RAM (GB) | DB loops | Runtime (s) | Throughput (reads/min) |
| --- | --- | --- | --- | --- | --- |
| 96 | 76 | 0.252 | 1 | 220.9 | 223,339 |
| 289 | 121 | 0.300 | 1 | 220.7 | 223,606 |
| 501 | 172 | 0.353 | 1 | 224.1 | 220,141 |

KENT achieved a mean precision of 58.55%, recall of 93.36%, and F1 score of 71.74%. CLARK-l produced highly similar results, with a mean precision of 58.33%, a recall of 93.42%, and an F1 score of 71.59%. The mean L1 abundance distance was 0.731 for KENT and 0.729 for CLARK-l. Thus, the source-level adaptations and distributed execution used by KENT did not produce a substantial change in species detection or abundance estimation performance relative to the CPU baseline.

The strict recall values also reflect the custom Marine reference database’s incomplete coverage of the CAMI2 community. Across the ten samples, species represented in the reference database accounted for 93.22% of the evaluated bacterial and archaeal ground-truth abundance. When recall was calculated only over ground-truth species represented in the reference database, CLARK-l recovered 99.93% *±* 0.15% and KENT recovered 99.86% of those species. These database-aware values are reported as a secondary analysis rather than as replacements for the conventional metrics in Table 3. They indicate that most of the false negatives in the strict evaluation arose from species absent from the reference database rather than from failure to detect reference-covered taxa.

### 3.4 Effect of Reference Database Size

To characterize the effect of reference database size on KENT performance, we evaluated the three nested CAMI2 Marine database subsets described in Section 2.5. After taxonomy mapping and removal of unidentified references, the databases contained 96, 289, and 501 unique mapped TaxIDs. All measurements were performed on the same Jetson Nano node using the same FASTQ fragment containing 822,406 reads and 128 CuCLARK-l input batches. Thus, the sequencing input and execution configuration were held fixed while the reference database was varied.

As the number of mapped TaxIDs increased from 96 to 501, the compiled database size on disk increased from 76 MB to 172 MB, while the database size reported by CuCLARK-l when loaded in RAM increased from 0.252 GB to 0.353 GB. All three configurations required a single CuCLARK-l database-processing loop. Classification runtimes were 220.9, 220.7, and 224.1 s for the 96-, 289-, and 501-TaxID databases, respectively, corresponding to throughputs of 223,339, 223,606, and 220,141 reads/min. Across the tested range, the reported in-memory database size increased by approximately 40%, whereas the runtime difference between the smallest and largest configurations was approximately 1.45%. No substantial degradation in classification throughput was observed while all three databases remained in single-loop configurations.

These results indicate that, within the tested range, KENT’s classification performance was relatively insensitive to reference database size when the database could be processed in a single CuCLARK-l database-processing loop. The Jetson Nano’s operational memory limit, however, was not determined solely by database size. All complete CAMI2 Marine samples, ranging from approximately 4.70 to 4.91 Gbp, required input fragmentation, whereas approximately half-sized fragments of 2.35 to 2.46 Gbp completed successfully. The practical memory boundary, therefore, depends on the combined memory requirements of the loaded database, the sequence data contained in each input batch, intermediate classification structures, and other system memory usage. We therefore report the CAMI2 observations as an empirical operating range rather than as a universal database-size or input-size threshold.

### 3.5 Power and Deployment Considerations

All experiments were conducted with the Jetson Nano modules configured in MAXN mode, corresponding to a nominal 10 W module-level power budget per device. For the complete 11-node KENT cluster, this corresponds to a nominal aggregate module budget of 110 W when all nodes are active. Importantly, these values describe the configured module power envelope and should not be interpreted as measured electrical consumption. Actual system-level consumption additionally depends on the carrier boards, storage devices, Ethernet infrastructure, cooling, and other peripherals.

Within this power-constrained embedded configuration, KENT achieved average speedups of approximately 3.1*×* for the wastewater dataset and 2.68*×* for the CAMI2 Marine dataset relative to the CPU-based CLARK-l baseline running on the same class of Jetson hardware. These results demonstrate that the GPU-accelerated workflow substantially reduces classification time while operating within the Jetson Nano’s embedded power configuration. Direct wall-power or energyper-sample measurements were not available in the present study; therefore, no claims are made regarding the measured energy efficiency.

## 4 Conclusion

In this study, we present KENT, a metagenomic read-classification framework designed for low-power, memory-constrained edge hardware. KENT adapts CuCLARK-l for execution on NVIDIA Jetson Nano modules and integrates the modified classifier with node-local orchestration, abundance estimation, split-run merging, report generation, and MPI-based sample-level coordination. Across the wastewater and CAMI2 Marine experiments, KENT provided substantial runtime improvements over the CPU-based CLARK-l baseline, with average speedups of approximately 3.1× and 2.68×, respectively, while maintaining closely aligned abundance profiles.

The CAMI2 Marine ground-truth analysis provides an independent evaluation of biological profiling performance beyond agreement with CLARK-l. Across ten long-read samples, KENT achieved a mean species-level precision of 58.55%, recall of 93.36%, F1 score of 71.74%, and L1 abundance distance of 0.731, closely matching the corresponding CLARK-l results. The custom Marine reference database covered 93.22% of the evaluated bacterial and archaeal ground-truth abundance on average, and KENT recovered 99.86% of ground-truth species represented in that database. These results indicate that the modifications required for embedded GPU execution preserve the taxonomic profiling behavior of the baseline while providing a substantial runtime advantage.

The controlled database-size experiment further showed that classification throughput remained largely stable as the CuCLARK-l in-memory database increased from 0.252 to 0.353 GB, with all tested configurations remaining within a single database-processing loop. Memory capacity nevertheless remains the principal limitation of the Jetson Nano platform. Complete CAMI2 Marine inputs of approximately 4.70 to 4.91 Gbp required fragmentation, whereas approximately 2.35 to 2.46 Gbp fragments were processed successfully. This indicates that the practical execution boundary depends on the combined memory requirements of the reference database, sequencing input, intermediate classifier structures, and system workload rather than on database size alone.

KENT therefore demonstrates the feasibility of local, offline metagenomic screening on inexpensive embedded GPU hardware while also exposing the memory constraints that must be considered in field deployment. Future work will focus on improved memory management and more automated handling of oversized sequencing inputs, as well as extending the framework to newer embedded GPU platforms with greater memory capacity.

## Acknowledgments

This work was supported by a grant from the Bridge to Türkiye Fund under the NVIDIA Applied Research Accelerator Program, and by the Turkish Institutes of Health (TÜSEB) Group-B R&D Project (TB–46712) awarded to C.A.

During the preparation of this manuscript, the authors used generative AI tools to edit, rewrite, and reorganize portions of the text; to assist with organizing, analyzing, and visualizing data; to generate and refine code used in the analysis; and to create illustrative images. No generative AI tools were used to create or alter primary research data, results, or scientific images. The authors manually reviewed and verified all outputs, including text, code, analyses, and figures, and take full responsibility for the accuracy and integrity of the content of this publication.

## Conflict of interest statement

C. Alkan is a co-founder of Lidya Genomics.

