## Supplementary material for "KENT: Fast Pathogen Screening with NVIDIA Jetson Cluster in Resource-Constrained Areas"

### A Appendix: Detailed Experimental Results

#### A1 Wastewater custom reference database

The following section lists the reference genomes used to create the database for the wastewater dataset. Abundance comparison tables are provided in section A6.

**Table A1:** Species included in the custom wastewater pathogen database (22 reference sequences). The compiled CuCLARK-l database occupied approximately 62 MB on disk and was reported as 0.237 GB in RAM at runtime.

| No. | Species / reference target |
|---|---|
| 1 | *Giardia intestinalis* |
| 2 | *Plasmodium falciparum* |
| 3 | *Escherichia coli K-12* |
| 4 | *Pseudomonas aeruginosa* |
| 5 | *Salmonella enterica* |
| 6 | *Clostridium tetani* |
| 7 | *Escherichia coli O157:H7* |
| 8 | *Orientia tsutsugamushi* |
| 9 | *Staphylococcus aureus* |
| 10 | *Coccidioides immitis* |
| 11 | *Mycobacterium tuberculosis* |
| 12 | *Listeria monocytogenes* |
| 13 | *Hepatitis A virus* |
| 14 | *Zika virus* |
| 15 | *Haemophilus influenzae* |
| 16 | *Streptococcus pneumoniae* |
| 17 | *Moraxella catarrhalis* |
| 18 | *Human orthopneumovirus* |
| 19 | *Vibrio cholerae* |
| 20 | *Acinetobacter baumannii* |
| 21 | *Helicobacter pylori* |
| 22 | *Enterobacter cloacae* |

#### A2 Wastewater dataset metadata

**Table A2:** Wastewater long read samples used for evaluation.

| Sample | Num. reads | Sum length (bp) | Min | Avg | Max |
|---|---|---|---|---|---|
| SRR7627523 | 739,468 | 5,399,845,105 | 92 | 7,302.3 | 48,662 |
| SRR7635124 | 461,649 | 4,735,781,782 | 138 | 10,258.4 | 95,136 |
| SRR7631476 | 378,859 | 4,486,221,055 | 142 | 11,841.4 | 109,176 |
| SRR7503035 | 839,634 | 4,220,888,228 | 26 | 5,027.1 | 52,113 |
| SRR7499264 | 720,067 | 3,342,624,183 | 18 | 4,642.1 | 54,416 |
| SRR7630418 | 703,509 | 3,231,034,980 | 54 | 4,592.7 | 46,566 |
| SRR7503007 | 522,729 | 2,943,366,029 | 106 | 5,630.8 | 45,432 |
| SRR7618093 | 469,003 | 2,942,881,293 | 109 | 6,274.8 | 73,476 |
| SRR7500591 | 586,119 | 2,825,097,351 | 77 | 4,820.0 | 33,536 |
| SRR7497945 | 494,663 | 2,489,331,291 | 114 | 5,032.4 | 21,514 |
| SRR7497167 | 627,655 | 2,444,715,951 | 131 | 3,895.0 | 31,456 |

#### A3 CAMI2 Marine dataset metadata

**Table A3:** CAMI2 Marine long read samples used for evaluation.

| Sample | Num. reads | Sum length (bp) | Min | Avg | Max |
|---|---|---|---|---|---|
| sample_0 | 1,644,811 | 4,874,039,793 | 100 | 2,963.3 | 11,938 |
| sample_1 | 1,641,591 | 4,871,611,007 | 100 | 2,967.6 | 11,938 |
| sample_2 | 1,656,266 | 4,914,486,507 | 100 | 2,967.2 | 11,938 |
| sample_3 | 1,648,299 | 4,879,109,718 | 100 | 2,960.1 | 11,938 |
| sample_4 | 1,585,522 | 4,702,118,599 | 100 | 2,965.7 | 11,938 |
| sample_5 | 1,632,786 | 4,849,577,905 | 100 | 2,970.1 | 11,938 |
| sample_6 | 1,634,731 | 4,849,854,642 | 100 | 2,966.8 | 11,938 |
| sample_7 | 1,612,220 | 4,782,829,598 | 100 | 2,966.6 | 11,938 |
| sample_8 | 1,637,443 | 4,854,540,378 | 100 | 2,964.7 | 11,938 |
| sample_9 | 1,617,260 | 4,795,417,306 | 100 | 2,965.1 | 11,938 |

#### A4 Wastewater runtime logs

**Table A4:** Wastewater classification runtimes for the Jetson-based modified CuCLARK pipeline and the CLARK-l baseline. Split samples were classified into two fragments, and their abundance outputs were merged afterward.

| Sample | System | Time (s) | Speed (reads/min) | Reads |
|---|---|---|---|---|
| SRR7627523 part 1 | KENT | 245.3 | 90429 | 369,734 |
| SRR7627523 part 2 | KENT | 245.7 | 90283 | 369,734 |
| SRR7627523 full | CLARK-l | 1259.8 | 35218 | 739,468 |
| SRR7635124 part 1 | KENT | 180.0 | 76935 | 230,825 |
| SRR7635124 part 2 | KENT | 178.7 | 77503 | 230,824 |
| SRR7635124 full | CLARK-l | 1191.9 | 23238 | 461,649 |
| SRR7631476 part 1 | KENT | 207.9 | 54660 | 189,430 |
| SRR7631476 part 2 | KENT | 207.6 | 54747 | 189,429 |
| SRR7631476 full | CLARK-l | 1164.9 | 19513 | 378,859 |
| SRR7503035 | KENT | 468.9 | 107430 | 839,634 |
| SRR7503035 | CLARK-l | 1114.6 | 45199 | 839,634 |
| SRR7499264 | KENT | 306.5 | 140954 | 720,067 |
| SRR7499264 | CLARK-l | 907.9 | 47587 | 720,067 |
| SRR7630418 | KENT | 294.6 | 143278 | 703,509 |
| SRR7630418 | CLARK-l | 885.3 | 47682 | 703,509 |
| SRR7503007 | KENT | 214.9 | 145923 | 522,729 |
| SRR7503007 | CLARK-l | 810.1 | 38713 | 522,729 |
| SRR7618093 | KENT | 269.7 | 104338 | 469,003 |
| SRR7618093 | CLARK-l | 807.0 | 34443 | 469,003 |
| SRR7500591 | KENT | 256.8 | 136944 | 586,119 |
| SRR7500591 | CLARK-l | 849.3 | 41405 | 586,119 |
| SRR7497945 | KENT | 180.5 | 164432 | 494,663 |
| SRR7497945 | CLARK-l | 744.86 | 39846 | 494,663 |
| SRR7497167 | KENT | 221.6 | 169960 | 627,655 |
| SRR7497167 | CLARK-l | 561.5 | 67067 | 627,655 |

#### A5 CAMI2 Marine long read runtime logs

**Table A5:** CAMI2 Marine long read classification runtimes. The source reference collection occupied approximately 2.8 GB on disk, while the compiled CuCLARK-l database occupied approximately 172 MB on disk and was reported as 0.353 GB in RAM at runtime. The number of input batches was increased to 128. Jetson-side runs were carried out on split input fragments.

| Sample | System | Time (s) | Speed (reads/min) | Reads |
|---|---|---|---|---|
| sample0 part 1 | KENT | 464.8 | 106156 | 822,406 |
| sample0 part 2 | KENT | 464.7 | 106186 | 822,405 |
| sample0 full | CLARK-l | 1532.2 | 64407 | 1,644,811 |
| sample1 part 1 | KENT | 464.4 | 106055 | 820,796 |
| sample1 part 2 | KENT | 464.8 | 105962 | 820,795 |
| sample1 full | CLARK-l | 1632.2 | 60346 | 1,641,591 |
| sample2 part 1 | KENT | 468.0 | 106159 | 828,133 |
| sample2 part 2 | KENT | 469.3 | 105875 | 828,133 |
| sample2 full | CLARK-l | 1568.3 | 63365 | 1,656,266 |
| sample3 part 1 | KENT | 228.2 | 216723 | 824,150 |
| sample3 part 2 | KENT | 226.4 | 218452 | 824,149 |
| sample3 full | CLARK-l | 1620.4 | 61034 | 1,648,299 |
| sample4 part 1 | KENT | 220.3 | 215895 | 792,761 |
| sample4 part 2 | KENT | 218.7 | 217525 | 792,761 |
| sample4 full | CLARK-l | 1623.5 | 58598 | 1,585,522 |
| sample5 part 1 | KENT | 226.7 | 216038 | 816,393 |
| sample5 part 2 | KENT | 224.9 | 217761 | 816,393 |
| sample5 full | CLARK-l | 1703.4 | 57512 | 1,632,786 |
| sample6 part 1 | KENT | 228.1 | 214996 | 817,366 |
| sample6 part 2 | KENT | 226.8 | 216237 | 817,365 |
| sample6 full | CLARK-l | 1707.6 | 57437 | 1,634,731 |
| sample7 part 1 | KENT | 223.0 | 216920 | 806,110 |
| sample7 part 2 | KENT | 224.0 | 215945 | 806,110 |
| sample7 full | CLARK-l | 1737.8 | 55663 | 1,612,220 |
| sample8 part 1 | KENT | 218.9 | 224383 | 818,722 |
| sample8 part 2 | KENT | 219.7 | 223607 | 818,721 |
| sample8 full | CLARK-l | 2057.9 | 47741 | 1,637,443 |
| sample9 part 1 | KENT | 457.1 | 106153 | 808,630 |
| sample9 part 2 | KENT | 457.9 | 105948 | 808,630 |
| sample9 full | CLARK-l | 1969.6 | 49266 | 1,617,260 |

#### A6 Wastewater Abundance Comparisons

This section provides abundance comparisons between KENT and CLARK-l baseline for the wastewater dataset. Each table reports, for every reference taxon in the custom pathogen database, the raw assigned read count and abundance measures produced by the two systems.

- **Count** denotes the number of reads assigned to a given taxon.
- **All (%)** denotes the percentage of all reads in the sample assigned to that taxon, including unclassified reads.
- **Class. (%)** denotes the percentage of only the classified reads assigned to that taxon, excluding unclassified reads.

**Table A6:** Taxon-level abundance comparison between KENT and CLARK-l for sample SRR7627523. Differences are reported as KENT minus CLARK-l.

| Name | KENT count | CLARK-l count | Count diff | KENT all (%) | CLARK-l all (%) | All diff | KENT class. (%) | CLARK-l class. (%) | Class. diff |
|---|---|---|---|---|---|---|---|---|---|
| Acinetobacter baumannii | 192 | 175 | -17 | 0.030000 | 0.023666 | -0.006334 | 0.420000 | 0.379610 | -0.040390 |
| Clostridium tetani | 62 | 82 | 20 | 0.010000 | 0.011089 | 0.001089 | 0.130000 | 0.177874 | 0.047874 |
| Coccidioides immitis | 20,554 | 20,309 | -245 | 2.780000 | 2.746430 | -0.033570 | 44.600000 | 44.054200 | -0.545800 |
| Enterobacter cloacae | 130 | 163 | 33 | 0.020000 | 0.022043 | 0.002043 | 0.280000 | 0.353579 | 0.073579 |
| Escherichia coli | 494 | 459 | -35 | 0.070000 | 0.062072 | -0.007928 | 1.070000 | 0.995662 | -0.074338 |
| Giardia intestinalis | 7 | 6 | -1 | 0.000000 | 0.000811 | 0.000811 | 0.020000 | 0.013015 | -0.006985 |
| Haemophilus influenzae | 224 | 164 | -60 | 0.030000 | 0.022178 | -0.007822 | 0.490000 | 0.355748 | -0.134252 |
| Helicobacter pylori | 15 | 20 | 5 | 0.000000 | 0.002705 | 0.002705 | 0.030000 | 0.043384 | 0.013384 |
| Listeria monocytogenes | 79 | 112 | 33 | 0.010000 | 0.015146 | 0.005146 | 0.170000 | 0.242950 | 0.072950 |
| Moraxella catarrhalis | 79 | 78 | -1 | 0.010000 | 0.010548 | 0.000548 | 0.170000 | 0.169197 | -0.000803 |
| Mycobacterium tuberculosis | 446 | 419 | -27 | 0.060000 | 0.056662 | -0.003338 | 0.970000 | 0.908894 | -0.061106 |
| Orientia tsutsugamushi | 2 | 2 | 0 | 0.000000 | 0.000270 | 0.000270 | 0.000000 | 0.004338 | 0.004338 |
| Plasmodium falciparum | 22,652 | 22,945 | 293 | 3.060000 | 3.102910 | 0.042910 | 49.150000 | 49.772200 | 0.622200 |
| Pseudomonas aeruginosa | 683 | 735 | 52 | 0.090000 | 0.099396 | 0.009396 | 1.480000 | 1.594360 | 0.114360 |
| Salmonella enterica | 274 | 188 | -86 | 0.040000 | 0.025424 | -0.014576 | 0.590000 | 0.407809 | -0.182191 |
| Staphylococcus aureus | 68 | 77 | 9 | 0.010000 | 0.010413 | 0.000413 | 0.150000 | 0.167028 | 0.017028 |
| Streptococcus pneumoniae | 8 | 15 | 7 | 0.000000 | 0.002028 | 0.002028 | 0.020000 | 0.032538 | 0.012538 |
| UNKNOWN | 693,380 | 693,368 | -12 | 93.770000 | 93.765800 | -0.004200 | – | – | – |
| Vibrio cholerae | 119 | 151 | 32 | 0.020000 | 0.020420 | 0.000420 | 0.260000 | 0.327549 | 0.067549 |

**Table A7:** Taxon-level abundance comparison between KENT and CLARK-l for sample SRR7635124. Differences are reported as KENT minus CLARK-l.

| Name | KENT count | CLARK-l count | Count diff | KENT all (%) | CLARK-l all (%) | All diff | KENT class. (%) | CLARK-l class. (%) | Class. diff |
|---|---|---|---|---|---|---|---|---|---|
| Acinetobacter baumannii | 128 | 129 | 1 | 0.030000 | 0.027943 | -0.002057 | 0.040000 | 0.038043 | -0.001957 |
| Clostridium tetani | 3 | 3 | 0 | 0.000000 | 0.000650 | 0.000650 | 0.000000 | 0.000885 | 0.000885 |
| Coccidioides immitis | 3,620 | 3,985 | 365 | 0.780000 | 0.863210 | 0.083210 | 1.090000 | 1.175220 | 0.085220 |
| Enterobacter cloacae | 7,157 | 6,959 | -198 | 1.550000 | 1.507420 | -0.042580 | 2.160000 | 2.052280 | -0.107720 |
| Escherichia coli | 309,342 | 318,137 | 8,795 | 67.010000 | 68.913200 | 1.903200 | 93.440000 | 93.821900 | 0.381900 |
| Giardia intestinalis | 1 | nan | nan | 0.000000 | – | – | 0.000000 | – | – |
| Haemophilus influenzae | 60 | 100 | 40 | 0.010000 | 0.021662 | 0.011661 | 0.020000 | 0.029491 | 0.009491 |
| Helicobacter pylori | 1 | 2 | 1 | 0.000000 | 0.000433 | 0.000433 | 0.000000 | 0.000590 | 0.000590 |
| Listeria monocytogenes | 7 | 4 | -3 | 0.000000 | 0.000866 | 0.000866 | 0.000000 | 0.001180 | 0.001180 |
| Moraxella catarrhalis | 18 | 10 | -8 | 0.000000 | 0.002166 | 0.002166 | 0.010000 | 0.002949 | -0.007051 |
| Mycobacterium tuberculosis | 6 | 6 | 0 | 0.000000 | 0.001300 | 0.001300 | 0.000000 | 0.001769 | 0.001769 |
| Orientia tsutsugamushi | nan | 1 | nan | – | 0.000217 | – | – | 0.000295 | – |
| Plasmodium falciparum | 4,145 | 4,131 | -14 | 0.900000 | 0.894836 | -0.005164 | 1.250000 | 1.218280 | -0.031720 |
| Pseudomonas aeruginosa | 380 | 503 | 123 | 0.080000 | 0.108957 | 0.028957 | 0.110000 | 0.148340 | 0.038340 |
| Salmonella enterica | 5,759 | 4,578 | -1,181 | 1.250000 | 0.991662 | -0.258338 | 1.740000 | 1.350100 | -0.389900 |
| Staphylococcus aureus | 12 | 10 | -2 | 0.000000 | 0.002166 | 0.002166 | 0.000000 | 0.002949 | 0.002949 |
| Streptococcus pneumoniae | 57 | 56 | -1 | 0.010000 | 0.012130 | 0.002130 | 0.020000 | 0.016515 | -0.003485 |
| UNKNOWN | 130,578 | 122,563 | -8,015 | 28.290000 | 26.549000 | -1.741000 | – | – | – |
| Vibrio cholerae | 375 | 472 | 97 | 0.080000 | 0.102242 | 0.022242 | 0.110000 | 0.139198 | 0.029198 |

**Table A8:** Taxon-level abundance comparison between KENT and CLARK-l for sample SRR7631476. Differences are reported as KENT minus CLARK-l.

| Name | KENT count | CLARK-l count | Count diff | KENT all (%) | CLARK-l all (%) | All diff | KENT class. (%) | CLARK-l class. (%) | Class. diff |
|---|---|---|---|---|---|---|---|---|---|
| Acinetobacter baumannii | 394 | 421 | 27 | 0.100000 | 0.111123 | 0.011123 | 0.170000 | 0.170567 | 0.000567 |
| Clostridium tetani | 12 | 12 | 0 | 0.000000 | 0.003167 | 0.003167 | 0.010000 | 0.004862 | -0.005138 |
| Coccidioides immitis | 113 | 126 | 13 | 0.030000 | 0.033258 | 0.003258 | 0.050000 | 0.051048 | 0.001048 |
| Enterobacter cloacae | 6,526 | 6,335 | -191 | 1.720000 | 1.672130 | -0.047870 | 2.760000 | 2.566610 | -0.193390 |
| Escherichia coli | 223,763 | 234,723 | 10,960 | 59.060000 | 61.955200 | 2.895200 | 94.500000 | 95.097300 | 0.597300 |
| Giardia intestinalis | 1 | nan | nan | 0.000000 | – | – | 0.000000 | – | – |
| Haemophilus influenzae | 60 | 98 | 38 | 0.020000 | 0.025867 | 0.005867 | 0.030000 | 0.039704 | 0.009704 |
| Helicobacter pylori | 5 | 14 | 9 | 0.000000 | 0.003695 | 0.003695 | 0.000000 | 0.005672 | 0.005672 |
| Listeria monocytogenes | 24 | 13 | -11 | 0.010000 | 0.003431 | -0.006569 | 0.010000 | 0.005267 | -0.004733 |
| Moraxella catarrhalis | 15 | 15 | 0 | 0.000000 | 0.003959 | 0.003959 | 0.010000 | 0.006077 | -0.003923 |
| Mycobacterium tuberculosis | 7 | 8 | 1 | 0.000000 | 0.002112 | 0.002112 | 0.000000 | 0.003241 | 0.003241 |
| Plasmodium falciparum | 174 | 184 | 10 | 0.050000 | 0.048567 | -0.001433 | 0.070000 | 0.074547 | 0.004547 |
| Pseudomonas aeruginosa | 201 | 269 | 68 | 0.050000 | 0.071003 | 0.021003 | 0.080000 | 0.108985 | 0.028985 |
| Salmonella enterica | 5,024 | 4,047 | -977 | 1.330000 | 1.068210 | -0.261790 | 2.120000 | 1.639630 | -0.480370 |
| Staphylococcus aureus | 43 | 39 | -4 | 0.010000 | 0.010294 | 0.000294 | 0.020000 | 0.015801 | -0.004199 |
| Streptococcus pneumoniae | 143 | 149 | 6 | 0.040000 | 0.039329 | -0.000671 | 0.060000 | 0.060367 | 0.000367 |
| UNKNOWN | 142,072 | 132,035 | -10,037 | 37.500000 | 34.850700 | -2.649300 | – | – | – |
| Vibrio cholerae | 282 | 371 | 89 | 0.070000 | 0.097926 | 0.027926 | 0.120000 | 0.150310 | 0.030310 |

**Table A9:** Taxon-level abundance comparison between KENT and CLARK-l for sample SRR7503035. Differences are reported as KENT minus CLARK-l.

| Name | KENT count | CLARK-l count | Count diff | KENT all (%) | CLARK-l all (%) | All diff | KENT class. (%) | CLARK-l class. (%) | Class. diff |
|---|---|---|---|---|---|---|---|---|---|
| Acinetobacter baumannii | 378 | 370 | -8 | 0.045020 | 0.044067 | -0.000953 | 0.900794 | 0.881099 | -0.019695 |
| Clostridium tetani | 99 | 113 | 14 | 0.011791 | 0.013458 | 0.001667 | 0.235922 | 0.269092 | 0.033170 |
| Coccidioides immitis | 15,809 | 16,028 | 219 | 1.882840 | 1.908930 | 0.026090 | 37.673700 | 38.168300 | 0.494600 |
| Enterobacter cloacae | 264 | 266 | 2 | 0.031442 | 0.031681 | 0.000238 | 0.629126 | 0.633439 | 0.004313 |
| Escherichia coli | 811 | 869 | 58 | 0.096590 | 0.103497 | 0.006907 | 1.932650 | 2.069390 | 0.136740 |
| Giardia intestinalis | 28 | 22 | -6 | 0.003335 | 0.002620 | -0.000715 | 0.066725 | 0.052390 | -0.014336 |
| Haemophilus influenzae | 130 | 138 | 8 | 0.015483 | 0.016436 | 0.000953 | 0.309797 | 0.328626 | 0.018829 |
| Helicobacter pylori | 18 | 29 | 11 | 0.002144 | 0.003454 | 0.001310 | 0.042895 | 0.069059 | 0.026164 |
| Listeria monocytogenes | 157 | 112 | -45 | 0.018699 | 0.013339 | -0.005359 | 0.374139 | 0.266711 | -0.107428 |
| Moraxella catarrhalis | 146 | 111 | -35 | 0.017389 | 0.013220 | -0.004169 | 0.347926 | 0.264330 | -0.083596 |
| Mycobacterium tuberculosis | 294 | 282 | -12 | 0.035015 | 0.033586 | -0.001429 | 0.700617 | 0.671540 | -0.029077 |
| Orientia tsutsugamushi | 15 | 16 | 1 | 0.001786 | 0.001906 | 0.000119 | 0.035746 | 0.038102 | 0.002356 |
| Plasmodium falciparum | 22,730 | 22,587 | -143 | 2.707130 | 2.690100 | -0.017030 | 54.166800 | 53.787500 | -0.379300 |
| Pseudomonas aeruginosa | 522 | 568 | 46 | 0.062170 | 0.067649 | 0.005479 | 1.243950 | 1.352610 | 0.108660 |
| Salmonella enterica | 323 | 198 | -125 | 0.038469 | 0.023582 | -0.014887 | 0.769726 | 0.471507 | -0.298219 |
| Staphylococcus aureus | 67 | 83 | 16 | 0.007980 | 0.009885 | 0.001906 | 0.159664 | 0.197652 | 0.037988 |
| Streptococcus pneumoniae | 11 | 14 | 3 | 0.001310 | 0.001667 | 0.000357 | 0.026214 | 0.033339 | 0.007125 |
| UNKNOWN | 797,671 | 797,641 | -30 | 95.002200 | 94.998700 | -0.003500 | – | – | – |
| Vibrio cholerae | 161 | 187 | 26 | 0.019175 | 0.022272 | 0.003097 | 0.383671 | 0.445312 | 0.061641 |

**Table A10:** Taxon-level abundance comparison between KENT and CLARK-l for sample SRR7499264. Differences are reported as KENT minus CLARK-l.

| Name | KENT count | CLARK-l count | Count diff | KENT all (%) | CLARK-l all (%) | All diff | KENT class. (%) | CLARK-l class. (%) | Class. diff |
|---|---|---|---|---|---|---|---|---|---|
| Acinetobacter baumannii | 139 | 168 | 29 | 0.019304 | 0.023331 | 0.004027 | 0.339629 | 0.409207 | 0.069578 |
| Clostridium tetani | 61 | 93 | 32 | 0.008471 | 0.012915 | 0.004444 | 0.149046 | 0.226525 | 0.077479 |
| Coccidioides immitis | 15,305 | 15,510 | 205 | 2.125500 | 2.153970 | 0.028470 | 37.395900 | 37.778600 | 0.382700 |
| Enterobacter cloacae | 187 | 180 | -7 | 0.025970 | 0.024998 | -0.000972 | 0.456911 | 0.438436 | -0.018475 |
| Escherichia coli | 574 | 650 | 76 | 0.079715 | 0.090269 | 0.010555 | 1.402500 | 1.583240 | 0.180740 |
| Giardia intestinalis | 20 | 32 | 12 | 0.002778 | 0.004444 | 0.001667 | 0.048868 | 0.077944 | 0.029077 |
| Haemophilus influenzae | 193 | 144 | -49 | 0.026803 | 0.019998 | -0.006805 | 0.471571 | 0.350749 | -0.120822 |
| Helicobacter pylori | 36 | 29 | -7 | 0.005000 | 0.004027 | -0.000972 | 0.087961 | 0.070637 | -0.017324 |
| Listeria monocytogenes | 103 | 104 | 1 | 0.014304 | 0.014443 | 0.000139 | 0.251668 | 0.253319 | 0.001651 |
| Moraxella catarrhalis | 145 | 95 | -50 | 0.020137 | 0.013193 | -0.006944 | 0.354289 | 0.231397 | -0.122892 |
| Mycobacterium tuberculosis | 910 | 927 | 17 | 0.126377 | 0.128738 | 0.002361 | 2.223470 | 2.257950 | 0.034480 |
| Orientia tsutsugamushi | 13 | 12 | -1 | 0.001805 | 0.001667 | -0.000139 | 0.031764 | 0.029229 | -0.002535 |
| Plasmodium falciparum | 22,367 | 22,291 | -76 | 3.106240 | 3.095680 | -0.010560 | 54.651000 | 54.295500 | -0.355500 |
| Pseudomonas aeruginosa | 307 | 334 | 27 | 0.042635 | 0.046385 | 0.003750 | 0.750116 | 0.813543 | 0.063427 |
| Salmonella enterica | 224 | 174 | -50 | 0.031108 | 0.024164 | -0.006944 | 0.547316 | 0.423822 | -0.123494 |
| Staphylococcus aureus | 50 | 85 | 35 | 0.006944 | 0.011805 | 0.004861 | 0.122169 | 0.207039 | 0.084870 |
| Streptococcus pneumoniae | 15 | 12 | -3 | 0.002083 | 0.001667 | -0.000417 | 0.036651 | 0.029229 | -0.007422 |
| UNKNOWN | 679,140 | 679,012 | -128 | 94.316200 | 94.298400 | -0.017800 | – | – | – |
| Vibrio cholerae | 278 | 215 | -63 | 0.038608 | 0.029858 | -0.008749 | 0.679258 | 0.523688 | -0.155570 |

**Table A11:** Taxon-level abundance comparison between KENT and CLARK-l for sample SRR7630418. Differences are reported as KENT minus CLARK-l.

| Name | KENT count | CLARK-l count | Count diff | KENT all (%) | CLARK-l all (%) | All diff | KENT class. (%) | CLARK-l class. (%) | Class. diff |
|---|---|---|---|---|---|---|---|---|---|
| Acinetobacter baumannii | 186 | 259 | 73 | 0.026439 | 0.036815 | 0.010377 | 0.592602 | 0.824447 | 0.231845 |
| Clostridium tetani | 94 | 117 | 23 | 0.013362 | 0.016631 | 0.003269 | 0.299487 | 0.372434 | 0.072947 |
| Coccidioides immitis | 11,389 | 11,481 | 92 | 1.618880 | 1.631960 | 0.013080 | 36.285700 | 36.546200 | 0.260500 |
| Enterobacter cloacae | 454 | 338 | -116 | 0.064534 | 0.048045 | -0.016489 | 1.446460 | 1.075920 | -0.370540 |
| Escherichia coli | 1,261 | 1,182 | -79 | 0.179244 | 0.168015 | -0.011229 | 4.017590 | 3.762530 | -0.255060 |
| Giardia intestinalis | 18 | 20 | 2 | 0.002559 | 0.002843 | 0.000284 | 0.057349 | 0.063664 | 0.006315 |
| Haemophilus influenzae | 177 | 218 | 41 | 0.025160 | 0.030988 | 0.005828 | 0.563928 | 0.693936 | 0.130008 |
| Helicobacter pylori | 20 | 34 | 14 | 0.002843 | 0.004833 | 0.001990 | 0.063721 | 0.108229 | 0.044508 |
| Listeria monocytogenes | 98 | 122 | 24 | 0.013930 | 0.017342 | 0.003411 | 0.312231 | 0.388350 | 0.076119 |
| Moraxella catarrhalis | 107 | 119 | 12 | 0.015210 | 0.016915 | 0.001706 | 0.340905 | 0.378800 | 0.037895 |
| Mycobacterium tuberculosis | 480 | 479 | -1 | 0.068229 | 0.068087 | -0.000142 | 1.529300 | 1.524750 | -0.004550 |
| Orientia tsutsugamushi | 11 | 9 | -2 | 0.001564 | 0.001279 | -0.000284 | 0.035046 | 0.028649 | -0.006398 |
| Plasmodium falciparum | 15,985 | 15,897 | -88 | 2.272180 | 2.259670 | -0.012510 | 50.928700 | 50.603200 | -0.325500 |
| Pseudomonas aeruginosa | 483 | 520 | 37 | 0.068656 | 0.073915 | 0.005259 | 1.538850 | 1.655260 | 0.116410 |
| Salmonella enterica | 195 | 222 | 27 | 0.027718 | 0.031556 | 0.003838 | 0.621276 | 0.706669 | 0.085393 |
| Staphylococcus aureus | 192 | 113 | -79 | 0.027292 | 0.016062 | -0.011229 | 0.611718 | 0.359701 | -0.252017 |
| Streptococcus pneumoniae | 14 | 20 | 6 | 0.001990 | 0.002843 | 0.000853 | 0.044604 | 0.063664 | 0.019059 |
| UNKNOWN | 672,122 | 672,094 | -28 | 95.538500 | 95.534500 | -0.004000 | – | – | – |
| Vibrio cholerae | 223 | 265 | 42 | 0.031698 | 0.037668 | 0.005970 | 0.710485 | 0.843546 | 0.133061 |

**Table A12:** Taxon-level abundance comparison between KENT and CLARK-l for sample SRR7503007. Differences are reported as KENT minus CLARK-l.

| Name | KENT count | CLARK-l count | Count diff | KENT all (%) | CLARK-l all (%) | All diff | KENT class. (%) | CLARK-l class. (%) | Class. diff |
|---|---|---|---|---|---|---|---|---|---|
| Acinetobacter baumannii | 93 | 93 | 0 | 0.017791 | 0.017791 | 0.000000 | 0.407341 | 0.407180 | -0.000161 |
| Clostridium tetani | 48 | 50 | 2 | 0.009183 | 0.009565 | 0.000383 | 0.210240 | 0.218914 | 0.008674 |
| Coccidioides immitis | 9,437 | 9,535 | 98 | 1.805330 | 1.824080 | 0.018750 | 41.334200 | 41.746900 | 0.412700 |
| Enterobacter cloacae | 89 | 71 | -18 | 0.017026 | 0.013583 | -0.003443 | 0.389821 | 0.310858 | -0.078963 |
| Escherichia coli | 286 | 324 | 38 | 0.054713 | 0.061982 | 0.007269 | 1.252680 | 1.418560 | 0.165880 |
| Giardia intestinalis | 7 | 4 | -3 | 0.001339 | 0.000765 | -0.000574 | 0.030660 | 0.017513 | -0.013147 |
| Haemophilus influenzae | 67 | 70 | 3 | 0.012817 | 0.013391 | 0.000574 | 0.293461 | 0.306480 | 0.013019 |
| Helicobacter pylori | 7 | 15 | 8 | 0.001339 | 0.002870 | 0.001530 | 0.030660 | 0.065674 | 0.035014 |
| Listeria monocytogenes | 83 | 61 | -22 | 0.015878 | 0.011669 | -0.004209 | 0.363541 | 0.267075 | -0.096466 |
| Moraxella catarrhalis | 55 | 44 | -11 | 0.010522 | 0.008417 | -0.002104 | 0.240901 | 0.192644 | -0.048257 |
| Mycobacterium tuberculosis | 90 | 87 | -3 | 0.017217 | 0.016643 | -0.000574 | 0.394201 | 0.380911 | -0.013290 |
| Orientia tsutsugamushi | 7 | 5 | -2 | 0.001339 | 0.000957 | -0.000383 | 0.030660 | 0.021891 | -0.008769 |
| Plasmodium falciparum | 11,888 | 11,815 | -73 | 2.274220 | 2.260250 | -0.013970 | 52.069600 | 51.729400 | -0.340200 |
| Pseudomonas aeruginosa | 400 | 423 | 23 | 0.076522 | 0.080921 | 0.004400 | 1.752000 | 1.852010 | 0.100010 |
| Salmonella enterica | 158 | 95 | -63 | 0.030226 | 0.018174 | -0.012052 | 0.692042 | 0.415937 | -0.276105 |
| Staphylococcus aureus | 36 | 49 | 13 | 0.006887 | 0.009374 | 0.002487 | 0.157680 | 0.214536 | 0.056856 |
| Streptococcus pneumoniae | 5 | 11 | 6 | 0.000957 | 0.002104 | 0.001148 | 0.021900 | 0.048161 | 0.026261 |
| UNKNOWN | 499,898 | 499,889 | -9 | 95.632300 | 95.630600 | -0.001700 | – | – | – |
| Vibrio cholerae | 75 | 88 | 13 | 0.014348 | 0.016835 | 0.002487 | 0.328501 | 0.385289 | 0.056788 |

**Table A13:** Taxon-level abundance comparison between KENT and CLARK-l for sample SRR7618093. Differences are reported as KENT minus CLARK-l.

| Name | KENT count | CLARK-l count | Count diff | KENT all (%) | CLARK-l all (%) | All diff | KENT class. (%) | CLARK-l class. (%) | Class. diff |
|---|---|---|---|---|---|---|---|---|---|
| Acinetobacter baumannii | 784 | 811 | 27 | 0.167163 | 0.172920 | 0.005757 | 2.096090 | 2.164740 | 0.068650 |
| Clostridium tetani | 123 | 126 | 3 | 0.026226 | 0.026866 | 0.000640 | 0.328851 | 0.336323 | 0.007472 |
| Coccidioides immitis | 12,017 | 11,858 | -159 | 2.562240 | 2.528340 | -0.033900 | 32.128400 | 31.651700 | -0.476700 |
| Enterobacter cloacae | 468 | 596 | 128 | 0.099786 | 0.127078 | 0.027292 | 1.251240 | 1.590860 | 0.339620 |
| Escherichia coli | 2,255 | 2,228 | -27 | 0.480807 | 0.475050 | -0.005757 | 6.028930 | 5.947040 | -0.081890 |
| Giardia intestinalis | 9 | 6 | -3 | 0.001919 | 0.001279 | -0.000640 | 0.024062 | 0.016015 | -0.008047 |
| Haemophilus influenzae | 235 | 171 | -64 | 0.050106 | 0.036460 | -0.013646 | 0.628292 | 0.456438 | -0.171854 |
| Helicobacter pylori | 79 | 129 | 50 | 0.016844 | 0.027505 | 0.010661 | 0.211213 | 0.344331 | 0.133118 |
| Listeria monocytogenes | 157 | 167 | 10 | 0.033475 | 0.035607 | 0.002132 | 0.419752 | 0.445761 | 0.026009 |
| Moraxella catarrhalis | 157 | 157 | 0 | 0.033475 | 0.033475 | 0.000000 | 0.419752 | 0.419069 | -0.000683 |
| Mycobacterium tuberculosis | 96 | 105 | 9 | 0.020469 | 0.022388 | 0.001919 | 0.256664 | 0.280269 | 0.023605 |
| Orientia tsutsugamushi | 3 | 2 | -1 | 0.000640 | 0.000426 | -0.000213 | 0.008021 | 0.005338 | -0.002682 |
| Plasmodium falciparum | 18,925 | 19,168 | 243 | 4.035160 | 4.086970 | 0.051810 | 50.597500 | 51.163800 | 0.566300 |
| Pseudomonas aeruginosa | 1,164 | 990 | -174 | 0.248186 | 0.211086 | -0.037100 | 3.112050 | 2.642540 | -0.469510 |
| Salmonella enterica | 504 | 384 | -120 | 0.107462 | 0.081876 | -0.025586 | 1.347490 | 1.024980 | -0.322510 |
| Staphylococcus aureus | 100 | 162 | 62 | 0.021322 | 0.034541 | 0.013220 | 0.267358 | 0.432415 | 0.165057 |
| Streptococcus pneumoniae | 45 | 34 | -11 | 0.009595 | 0.007249 | -0.002345 | 0.120311 | 0.090754 | -0.029557 |
| UNKNOWN | 431,600 | 431,539 | -61 | 92.025000 | 92.012000 | -0.013000 | – | – | – |
| Vibrio cholerae | 282 | 370 | 88 | 0.060128 | 0.078891 | 0.018763 | 0.753950 | 0.987615 | 0.233665 |

**Table A14:** Taxon-level abundance comparison between KENT and CLARK-l for sample SRR7500591. Differences are reported as KENT minus CLARK-l.

| Name | KENT count | CLARK-l count | Count diff | KENT all (%) | CLARK-l all (%) | All diff | KENT class. (%) | CLARK-l class. (%) | Class. diff |
|---|---|---|---|---|---|---|---|---|---|
| Acinetobacter baumannii | 531 | 555 | 24 | 0.090596 | 0.094691 | 0.004095 | 1.343660 | 1.404390 | 0.060730 |
| Clostridium tetani | 311 | 307 | -4 | 0.053061 | 0.052378 | -0.000682 | 0.786963 | 0.776842 | -0.010121 |
| Coccidioides immitis | 12,479 | 12,243 | -236 | 2.129090 | 2.088820 | -0.040270 | 31.577200 | 30.980000 | -0.597200 |
| Enterobacter cloacae | 633 | 664 | 31 | 0.107999 | 0.113288 | 0.005289 | 1.601760 | 1.680200 | 0.078440 |
| Escherichia coli | 2,460 | 2,178 | -282 | 0.419710 | 0.371597 | -0.048113 | 6.224850 | 5.511270 | -0.713580 |
| Giardia intestinalis | 11 | 9 | -2 | 0.001877 | 0.001536 | -0.000341 | 0.027835 | 0.022774 | -0.005061 |
| Haemophilus influenzae | 158 | 213 | 55 | 0.026957 | 0.036341 | 0.009384 | 0.399808 | 0.538981 | 0.139173 |
| Helicobacter pylori | 72 | 102 | 30 | 0.012284 | 0.017403 | 0.005118 | 0.182191 | 0.258104 | 0.075913 |
| Listeria monocytogenes | 302 | 254 | -48 | 0.051525 | 0.043336 | -0.008190 | 0.764189 | 0.642729 | -0.121460 |
| Moraxella catarrhalis | 195 | 109 | -86 | 0.033270 | 0.018597 | -0.014673 | 0.493434 | 0.275817 | -0.217617 |
| Mycobacterium tuberculosis | 107 | 122 | 15 | 0.018256 | 0.020815 | 0.002559 | 0.270756 | 0.308712 | 0.037956 |
| Orientia tsutsugamushi | 7 | 8 | 1 | 0.001194 | 0.001365 | 0.000171 | 0.017713 | 0.020243 | 0.002530 |
| Plasmodium falciparum | 21,077 | 21,334 | 257 | 3.596030 | 3.639880 | 0.043850 | 53.333800 | 53.984200 | 0.650400 |
| Pseudomonas aeruginosa | 382 | 453 | 71 | 0.065174 | 0.077288 | 0.012114 | 0.966624 | 1.146280 | 0.179656 |
| Salmonella enterica | 273 | 342 | 69 | 0.046578 | 0.058350 | 0.011772 | 0.690807 | 0.865407 | 0.174600 |
| Staphylococcus aureus | 193 | 266 | 73 | 0.032928 | 0.045383 | 0.012455 | 0.488373 | 0.673094 | 0.184721 |
| Streptococcus pneumoniae | 102 | 107 | 5 | 0.017403 | 0.018256 | 0.000853 | 0.258104 | 0.270756 | 0.012652 |
| UNKNOWN | 546,600 | 546,600 | 0 | 93.257500 | 93.257500 | 0.000000 | – | – | – |
| Vibrio cholerae | 226 | 253 | 27 | 0.038559 | 0.043165 | 0.004607 | 0.571877 | 0.640198 | 0.068321 |

**Table A15:** Taxon-level abundance comparison between KENT and CLARK-l for sample SRR7497945. Differences are reported as KENT minus CLARK-l.

| Name | KENT count | CLARK-l count | Count diff | KENT all (%) | CLARK-l all (%) | All diff | KENT class. (%) | CLARK-l class. (%) | Class. diff |
|---|---|---|---|---|---|---|---|---|---|
| Acinetobacter baumannii | 182 | 173 | -9 | 0.036793 | 0.034973 | -0.001819 | 0.559432 | 0.531768 | -0.027664 |
| Clostridium tetani | 68 | 73 | 5 | 0.013747 | 0.014757 | 0.001011 | 0.209019 | 0.224388 | 0.015369 |
| Coccidioides immitis | 9,670 | 9,758 | 88 | 1.954870 | 1.972660 | 0.017790 | 29.723700 | 29.994200 | 0.270500 |
| Enterobacter cloacae | 263 | 233 | -30 | 0.053167 | 0.047103 | -0.006065 | 0.808410 | 0.716196 | -0.092214 |
| Escherichia coli | 7,217 | 7,264 | 47 | 1.458970 | 1.468470 | 0.009500 | 22.183600 | 22.328100 | 0.144500 |
| Giardia intestinalis | 9 | 9 | 0 | 0.001819 | 0.001819 | 0.000000 | 0.027664 | 0.027664 | 0.000000 |
| Haemophilus influenzae | 234 | 255 | 21 | 0.047305 | 0.051550 | 0.004245 | 0.719270 | 0.783820 | 0.064550 |
| Helicobacter pylori | 68 | 83 | 15 | 0.013747 | 0.016779 | 0.003032 | 0.209019 | 0.255126 | 0.046107 |
| Listeria monocytogenes | 162 | 123 | -39 | 0.032750 | 0.024865 | -0.007884 | 0.497956 | 0.378078 | -0.119878 |
| Moraxella catarrhalis | 151 | 118 | -33 | 0.030526 | 0.023855 | -0.006671 | 0.464144 | 0.362709 | -0.101435 |
| Mycobacterium tuberculosis | 137 | 133 | -4 | 0.027696 | 0.026887 | -0.000809 | 0.421111 | 0.408816 | -0.012295 |
| Orientia tsutsugamushi | 5 | 5 | 0 | 0.001011 | 0.001011 | 0.000000 | 0.015369 | 0.015369 | 0.000000 |
| Plasmodium falciparum | 12,886 | 12,815 | -71 | 2.605010 | 2.590650 | -0.014360 | 39.609000 | 39.390800 | -0.218200 |
| Pseudomonas aeruginosa | 706 | 767 | 61 | 0.142723 | 0.155055 | 0.012332 | 2.170100 | 2.357610 | 0.187510 |
| Salmonella enterica | 329 | 212 | -117 | 0.066510 | 0.042857 | -0.023652 | 1.011280 | 0.651646 | -0.359634 |
| Staphylococcus aureus | 73 | 89 | 16 | 0.014757 | 0.017992 | 0.003234 | 0.224388 | 0.273568 | 0.049180 |
| Streptococcus pneumoniae | 24 | 29 | 5 | 0.004852 | 0.005863 | 0.001011 | 0.073771 | 0.089140 | 0.015369 |
| UNKNOWN | 462,130 | 462,130 | 0 | 93.423200 | 93.423200 | 0.000000 | – | – | – |
| Vibrio cholerae | 349 | 394 | 45 | 0.070553 | 0.079650 | 0.009097 | 1.072760 | 1.211080 | 0.138320 |

**Table A16:** Taxon-level abundance comparison between KENT and CLARK-l for sample SRR7497167. Differences are reported as KENT minus CLARK-l.

| Name | KENT count | CLARK-l count | Count diff | KENT all (%) | CLARK-l all (%) | All diff | KENT class. (%) | CLARK-l class. (%) | Class. diff |
|---|---|---|---|---|---|---|---|---|---|
| Acinetobacter baumannii | 683 | 670 | -13 | 0.108818 | 0.106747 | -0.002071 | 2.081430 | 2.041810 | -0.039620 |
| Clostridium tetani | 169 | 159 | -10 | 0.026926 | 0.025332 | -0.001593 | 0.515024 | 0.484549 | -0.030475 |
| Coccidioides immitis | 9,583 | 9,696 | 113 | 1.526790 | 1.544800 | 0.018010 | 29.204000 | 29.548400 | 0.344400 |
| Enterobacter cloacae | 527 | 490 | -37 | 0.083963 | 0.078068 | -0.005895 | 1.606020 | 1.493270 | -0.112750 |
| Escherichia coli | 2,028 | 2,103 | 75 | 0.323107 | 0.335057 | 0.011950 | 6.180290 | 6.408850 | 0.228560 |
| Giardia intestinalis | 10 | 9 | -1 | 0.001593 | 0.001434 | -0.000159 | 0.030475 | 0.027427 | -0.003047 |
| Haemophilus influenzae | 126 | 179 | 53 | 0.020075 | 0.028519 | 0.008444 | 0.383982 | 0.545499 | 0.161517 |
| Helicobacter pylori | 129 | 170 | 41 | 0.020553 | 0.027085 | 0.006532 | 0.393125 | 0.518072 | 0.124947 |
| Listeria monocytogenes | 311 | 243 | -68 | 0.049550 | 0.038715 | -0.010834 | 0.947766 | 0.740538 | -0.207228 |
| Moraxella catarrhalis | 178 | 126 | -52 | 0.028359 | 0.020075 | -0.008285 | 0.542451 | 0.383982 | -0.158469 |
| Mycobacterium tuberculosis | 212 | 209 | -3 | 0.033777 | 0.033299 | -0.000478 | 0.646066 | 0.636923 | -0.009143 |
| Orientia tsutsugamushi | 4 | 2 | -2 | 0.000637 | 0.000319 | -0.000319 | 0.012190 | 0.006095 | -0.006095 |
| Plasmodium falciparum | 15,871 | 15,780 | -91 | 2.528620 | 2.514120 | -0.014500 | 48.366600 | 48.089200 | -0.277400 |
| Pseudomonas aeruginosa | 588 | 655 | 67 | 0.093682 | 0.104357 | 0.010675 | 1.791920 | 1.996100 | 0.204180 |
| Salmonella enterica | 482 | 293 | -189 | 0.076794 | 0.046682 | -0.030112 | 1.468890 | 0.892912 | -0.575978 |
| Staphylococcus aureus | 170 | 205 | 35 | 0.027085 | 0.032661 | 0.005576 | 0.518072 | 0.624733 | 0.106661 |
| Streptococcus pneumoniae | 77 | 92 | 15 | 0.012268 | 0.014658 | 0.002390 | 0.234656 | 0.280368 | 0.045712 |
| UNKNOWN | 594,841 | 594,841 | 0 | 94.772000 | 94.772000 | 0.000000 | – | – | – |
| Vibrio cholerae | 1,666 | 1,733 | 67 | 0.265432 | 0.276107 | 0.010675 | 5.077100 | 5.281280 | 0.204180 |

#### A7 Implementation Differences Between CuCLARK and KENT

**Table A17:** Implementation-level comparison between the original CuCLARK software and KENT

| Component | Original CuCLARK | KENT |
|---|---|---|
| CUDA target | `sm_30` | `sm_53` for Jetson Nano |
| Classifier | CuCLARK or CuCLARK-l | Modified CuCLARK-l |
| Input batching | User-defined input batches | Batch count tuned for Jetson memory |
| Free-memory threshold | 1 GB | 200 MB |
| Reserved memory (light mode) | 300 MB | 150 MB |
| Execution model | Single host, one or more GPUs | Sample-level MPI execution across Jetsons |
| Orchestration | Scripts and command-line tools | Node-local `kent` workflow |
| Abundance estimation | External CLARK-compatible post-processing | Integrated into the `kent` workflow |
| Fragmented inputs | No integrated merging | Count-based abundance merging |
| Distributed coordination | Not supported | Configuration, synchronization, and status via `kent-mpi` |
| Outputs | CLARK-compatible classifications | Classifications, abundances, and run summaries |

#### A8 CAMI2 Marine Abundance Comparisons

We do not share the abundance comparison tables for the CAMI2 Marine dataset here since they are quite large. You can find them in the project repository: https://github.com/BilkentCompGen/kent
